# (*2R,6R*)-hydroxynorketamine facilitates extinction and prevents emotional impairment and stress-induced reinstatement in morphine abstinent mice

**DOI:** 10.1101/2023.12.07.570550

**Authors:** Andria Michael, Anna Onisiforou, Polymnia Georgiou, Morfeas Koumas, Elmar Mammadov, Panos Zanos

## Abstract

Opioid addiction is a pressing public health concern marked by frequent relapse during periods of abstinence, perpetuated by negative affective states and anhedonia-driven behaviors. In addition to the current epidemic that was declared in the U.S.A., opioid-related deaths are increasing in other countries around the world. Classical antidepressants, or the currently prescribed opioid substitution pharmacotherapies have limited efficacy to reverse maladaptive behavioral responses, negative affect or prevent relapse in opioid abstinent individuals. Here, by establishing and using novel mouse models for the study of opioid addiction, we demonstrate, for the first time, the therapeutic potential of ketamine’s metabolite, (*2R,6R*)-hydroxynorketamine (HNK). In particular, our studies showcase (*2R,6R*)-HNK’s ability to reverse conditioning to sub-effective doses of morphine in stress-susceptible mice, prevent conditioned-place aversion and mitigate acute somatic withdrawal symptoms in opioid-dependent animals. In addition, we show that this metabolite reverses anhedonia, anxiety-like behaviors, cognitive impairment, and general stress susceptibility associated with protracted opioid withdrawal, thereby presenting a promising therapeutic avenue for opioid relapse prevention. Our results strongly suggest that (*2R,6R*)-HNK, potentially by augmenting downstream brain-derived neurotrophic factor (BDNF) and GluN2A *N*-methyl-D-aspartate receptor signaling, effectively reverses maladaptive behavioral responses typical of protracted opioid abstinence. Furthermore, it facilitates the extinction of opioid conditioning and prevents stress-induced reinstatement of opioid-seeking behaviors. Our findings highlight how (*2R,6R*)-HNK, through an enhancement of synaptic plasticity in mood-regulating brain areas, has the potential to be an effective, next-generation pharmacotherapy for opioid use disorders by addressing emotional disturbances associated with protracted abstinence.

**SIGNIFICANCE STATEMENT:** Our studies represent a comprehensive exploration into the critical facets of opioid addiction and abstinence, offering critical insights into the significant potential of (*2R,6R*)-HNK as a treatment for this disorder. By unraveling the complex dynamics of opioid withdrawal and addressing the profound emotional disturbances underlying relapse vulnerability, our findings illuminate a promising innovative avenue for therapeutic intervention. The demonstrated ability of (*2R,6R*)-HNK to reverse maladaptive behaviors emerging during protracted opioid abstinence, facilitate extinction of opioid conditioning, prevent stress-induced reinstatement, represents a paradigm shift in addiction research. These revelations not only deepen our comprehension of the neurobehavioral complexities associated with opioid abstinence but also underscore the profound implications of (*2R,6R*)-HNK as a prospective pharmacotherapy in mitigating the devastating impact of opioid use disorder, potentially transforming addiction treatment strategies.

## INTRODUCTION

Opioid addiction, also known as opioid use disorders (OUDs), is a brain disorder characterized by a progressive impairment of the initially engaged brain reward system. This leads to a subsequent recruitment of aversion-regulating brain circuitries during periods of abstinence, characterized by enhanced sensitivity to stress, and increased risk for relapse ^1^. OUD is considered a chronic, relapsing brain disorder causing high rates of mortality, primarily due to overdose incidences, and it was declared a national emergency and an epidemic in the USA in 2017 ^2^; however, opioid-related deaths are also on the rise in Europe and other countries of the world ^3^. Prolonged abstinence from chronic opioid abuse is associated with lowered mood and reduced motivation for natural rewards (i.e., anhedonia; ^4^), increased anxiety ^5–7^, and social avoidance ^8^. The negative affect emerging during protracted abstinence from opioid abuse resembles some of the symptomatology observed in mood disorders ^7,9^. This aspect constitutes a major consideration for the effective treatment of OUDs, as it causes longer illness duration, poorer prognosis and higher relapse rates ^1,4^. In addition to this negative affect characterizing protracted opioid abstinence, exposure to stress during that period of abstinence is amongst the most critical factors that triggers relapse in opioid addicts ^10^. Classical, monoamine-based antidepressant medications ^11^ or the currently used opioid substitution pharmacotherapies ^12,13^ have been shown to have limited efficacy to reverse the negative affect or prevent relapse in opioid abstinent individuals. Therefore, identifying novel effective medications for the treatment of opioid abstinence-induced negative affect is critical for the long-term management of OUDs and prevention of relapse.

Evidence suggests that the anesthetic drug ketamine, an *N*-methyl-D-aspartate receptor (NMDAR) antagonist, established as the prototype rapid-acting antidepressant ^14^, holds potential for the treatment of certain facets of opioid addiction, including maintenance of opioid abstinence. In a double-blind, non-placebo-controlled clinical trial, intramuscular (i.m.) administration of ketamine (0.2 *vs* 2 mg/kg) in heroin abstinent individuals induced a dose-dependent enhancement of abstinence rates and longer-lasting decrease in heroin craving, with higher doses of ketamine being more effective ^15^. The same authors also compared the effectiveness of a single *vs* three consecutive ketamine sessions (2 mg/kg, i.m.) to maintain abstinence in detoxified heroin dependent patients, showing that repeated administration of ketamine was superior in prolonging abstinence compared with the single administration group ^16^. Moreover, in a randomized double-blind, placebo-controlled study conducted in opioid-dependent patients, pre-treatment with a sub-anesthetic dose of ketamine (0.5 mg/kg/h, i.v.) reduced precipitated opioid withdrawal symptoms following administration of an opioid receptor antagonist under general anesthesia ^17^. However, in this study there was no long-term beneficial effect of ketamine, since these patients reported no significant improvements of their social life, no enhancement of abstinence rates or treatment retention at four months after ketamine administration ^17^. Although these studies offer some evidence suggesting the potential use of ketamine in assisting with maintenance of opioid abstinence, these studies had certain limitations. Two of these studies lacked inactive placebo control groups and the third study only assessed the effect of ketamine following acute precipitated withdrawal by using an opioid receptor antagonist under general anesthesia.

Similar to the clinical findings, racemic ketamine administration reduced somatic withdrawal symptoms following administration of an opioid receptor antagonist in morphine-dependent rats ^18^. In line with this finding, a more recent preclinical trial showed that the (*R*)-ketamine enantiomer can also reduce acute physical withdrawal symptoms precipitated by naloxone in oxycodone-dependent rats ^19^, while the (*S*)-ketamine enantiomer was not tested. However, ketamine’s potential to be used as an everyday treatment for opioid addiction is limited due to its side effects including dissociation and abuse potential ^20^. Ketamine, a racemic mixture consisting of the (*S*)- and (*R*)-ketamine enantiomers, is rapidly and stereoselectively metabolized to a number of metabolites, including *N*-demethylated norketamines and dehydronorketamines, as well as hydroxynorketamines (HNKs) ^21^. We have previously provided evidence that (*2R,6R*)-hydroxynorketamine (HNK), similar to its parent drug ketamine, rapidly reverses negative affective behaviors via a presynaptic mechanism converging with the mGlu_2_ receptor signaling to enhance excitatory neurotransmission, which subsequently increases brain-derived neurotrophic factor (BDNF) levels and activates downstream postsynaptic mammalian target of rapamycin complex 1 (mTORC1) pathways and increases protein synthesis in the form of α-amino-3-hydroxy-5-methyl-4-isoxazolepropionic acid receptor (AMPAR)-mediated synaptic strengthening ^22–28^.

There are is some recent evidence showing that the (*2R,6R*)-HNK metabolite is able to reduce acute withdrawal precipitated somatic symptoms, and prevents priming-induced reinstatement of opioid-seeking behaviors following extinction ^29^. Nevertheless, there is no prior study assessing the efficacy of (*2R,6R*)-HNK in reversing the negative affect induced by protracted opioid withdrawal, nor stress-induced reinstatement.

In the present study, we have developed several novel mouse models for testing different behaviors associated with opioid addiction and in some experiments, we incorporated models assessing comorbid maladaptive behavioral responses. By using these innovative models, our aim was to determine whether ketamine’s metabolite (*2R,6R*)-HNK could serve as a potential treatment at various stages within the opioid addiction cycle.

## MATERIALS AND METHODS

### Animals

Male C57BL/6J mice (8 weeks old, Jackson laboratories) were kept in a temperature and humidity-controlled environment and maintained in a 12:12 hr light/dark cycle (lights on at 7:00AM). Mice were singly housed for all the studies described due to increased aggression induced by opioid administration and were randomly assigned to treatment groups. All experimental procedures were approved by the University of Maryland, Baltimore Animal Care and Use Committee and the Cyprus National Committee for Animal Welfare and were conducted in full accordance with the National Institutes of Health (NIH) Guide for the Care and Use of Laboratory Animals and reported according to *ARRIVE* guidelines.

### Drugs

(*2R,6R*)-HNK hydrochloride was synthesized and characterized at the National Center for Advancing Translational Sciences. Absolute and relative stereochemistry for (*2R,6R*)-HNK was confirmed by small molecule x-ray crystallography ^27^. Morphine hydrochloride was purchased from Cayman Europe and was injected subcutaneously (s.c.) at a volume of 4 ml/kg, or as otherwise stated in the Methods section. Naloxone (Tocris, USA) was administered intraperitoneally (i.p.), at the dose of 1 mg/kg. Quinine sulfate dihydrate was purchased by Arcos Organics, USA. For the intraperitoneal injections, drugs were dissolved in 0.9% saline and given in a volume of 4 ml/kg.

#### Experiment 1: Effect of (*2R,6R*)-HNK on the development of conditioning to sub-effective doses of morphine in stress susceptible mice

To better understand the interplay between stress susceptibility and opioid dependence, we used an experimental design where mice were initially screened for stress susceptibility to develop maladaptive behaviors. We then assessed these mice for their propensity to develop conditioning to sub-effective doses of morphine. This experimental approach was conducted to establish, for the first time, a reliable animal model that mirrors the human scenario, where stress-susceptible individuals exhibit a heightened vulnerability to developing opioid dependence. We then assessed the efficacy of (*2R,6R*)-HNK to prevent conditioning to morphine in the stress susceptible mice.

### Chronic Social defeats

Socially defeated mice underwent a 10-day chronic social defeat stress paradigm, as described elsewhere ^30^. Briefly, experimental mice were introduced to the home cage (43 cm length x 11 cm width x 20 cm height) of a resident aggressive retired CD-1 breeder - prescreened for aggressive behaviors - for 10 min. Following this physical attack phase, mice were transferred and housed in the opposite side of the resident’s cage divided by a perforated Plexiglas divider, in order to maintain continuous, 24 h, sensory contact. This process was repeated daily for 10 days, with experimental mice being introduced to a novel aggressive CD-1 mouse each day.

### Social interaction test

Following the social defeats, on day 11, test mice were screened for susceptibility in a social interaction/avoidance choice test. The social interaction apparatus consisted of a rectangular three-chambered box (Stoelting Co., Wood Dale, IL, USA) comprised of two equal sized end-chambers and a smaller central chamber. The social interaction/avoidance choice test consisted of two 5-min phases. During the habituation phase, mice explored the empty apparatus. During the test phase, two small wire cages (Galaxy Cup, Spectrum Diversified Designs, Inc., Streetsboro, OH, USA), one containing a “stranger” CD-1 mouse and the other one empty, were placed in the far corners of each chamber. The time spent interacting (nose within close proximity to the cage) with the “stranger” mouse *versus* the empty cage was analyzed using TopScan video tracking software (CleverSys, Reston, Virginia). Mice with %Social Interaction ((Time spent interacting with stranger / total time interacting with the end-chamber cages) x 100) greater than 50% were considered as resilient, whereas mice with <50% %Social Interaction were considered as susceptible.

### Sucrose preference

Following the social interaction test mice were assessed for sucrose preference in their home cages overnight. Briefly, mice were singly housed and presented with two identical bottles containing either tap water or 1% (w/v) sucrose solution. Sucrose preference was measured (day 12) and were assigned to two groups: resilient (sucrose preference >70%) and susceptible (sucrose preference <55%).

### Final determination of stress susceptibility

Only mice that showed susceptibility and resilience in both the social interaction and sucrose preference test continued with the rest of the experimental procedures. Out of 98 total mice tested, only 6 mice did not meet the criteria for susceptibility in both the social interaction and sucrose preference tests and were excluded. 44 mice were assigned as susceptible and 48 mice as resilient.

### Test for sucrose preference deficits reversal

To avoid having the effects of acute stress affecting the opioid conditioning outcomes, we let mice recover from chronic social defeats stress for 20 days and re-tested for sucrose preference in their home cages (Day 30). Only mice that recovered normal sucrose preference (>70%) were used in the conditioning phase of the experiment. Only 3 mice that were initially assigned as resilient reverted sucrose preference to <70% and thus were excluded from the study. Therefore, 44 initially susceptible mice and 45 initially resilient mice were used in the morphine conditioning studies.

### Sub-effective dose morphine conditioning

The conditioned-place preference (CPP) apparatus consisted of a rectangular three-chambered box (40 cm length x 30 cm width x 35 cm height; Stoelting Co., Wood Dale, IL, USA) comprised of two equal sized end-chambers (20 cm x 18 cm x 35 cm) and a central chamber (20 cm x 10 cm x 35 cm). One end-chamber had a perforated floor and plain black walls, whereas the other end-chamber had a smooth floor and walls with vertical black and white stripes, as we previously described ^25^. The CPP protocol consisted of a pre-conditioning phase, eight conditioning sessions (over 4 days) and a post-conditioning test. On Day 32 (pre-conditioning phase), stress susceptible and resilient mice (that had successfully recovered from their sucrose preference deficits) were placed in the CPP apparatus and were allowed to explore all compartments for a period of 20 min. Drug-paired compartment was assigned in a non-biased and randomized way. On Day 33, mice were given a single injection of saline or (*2R,6R*)-HNK (10 mg/kg), to assess for possible effects of HNK to prevent stress-susceptibility-induced enhancement of morphine CPP.

### Conditioning phase (Days 34-37)

During the morning sessions of the conditioning phase, aline, or morphine at the sub-effective dose of 0.25 mg/kg were administered s.c., and mice were placed in their assigned compartment for 30 min. Five hrs later (afternoon sessions) saline was administered, and mice were placed in their drug-paired compartment for 30 min. During the post-conditioning test session (i.e. Day 38), mice were placed in the CPP apparatus to freely explore all three compartments for 20 min. Time spent in each compartment was measured during both pre- and post-conditioning sessions using TopScan v2.0 (CleverSys, Inc, VA, USA).

#### Experiment 2: Effect of (*2R,6R*)-HNK on the development of conditioning to high doses of morphine

We then determined whether the ability of (*2R,6R*)-HNK to prevent conditioning to sub-effective doses of morphine in stress susceptible mice (Experiment 1) was a result of its actions to reverse morphine conditioning *per se*, or a general action of this metabolite to reverse propensity to initiate conditioning to very low doses of morphine in mice having intrinsic stress susceptibility. In particular, we assessed (*2R,6R*)-HNK’s effects on high-dose morphine-induced conditioned-place preference in stress-naïve mice.

### Morphine conditioned-place preference

Here we used the same apparatus and protocol as used in Experiment 1. On Day 1 (pre-conditioning phase), mice were placed in the CPP apparatus and were allowed to explore all compartments for a period of 20 min. Drug-paired compartment was assigned in a non-biased and random way. On Day 2, mice were given a single injection of saline or (*2R,6R*)-HNK (10 mg/kg), to assess for possible effects of HNK to prevent morphine CPP. The conditioning phase (Days 3-6) was similar to the experiment above, but instead of 0.25 mg/kg morphine, we administered 5 mg/kg (s.c.) of the drug during the morning sessions. The post-conditioning test session was performed at Day 7. Time spent in each compartment was measured during both pre- and post-conditioning sessions using TopScan v2.0 (CleverSys, Inc, VA, USA).

#### Experiment 3: Effect of (*2R,6R*)-HNK on conditioned-place aversion induced by precipitated morphine withdrawal in opioid-dependent mice

To assess the potential therapeutic efficacy of (*2R,6R*)-HNK on the negative affective states associated with acute opioid abstinence, we made mice dependent to morphine and administered the opioid receptor antagonist naloxone to induce precipitated withdrawal. We investigated the ability of (*2R,6R*)-HNK to prevent conditioned-place aversion induced by precipitated morphine withdrawal in opioid-dependent mice.

### Precipitated morphine place aversion

The conditioned-place aversion (CPA) protocol was performed in a rectangular three-chambered box (40 cm length x 30 cm width x 35 cm height; Stoelting Co., Wood Dale, IL, USA) comprised of two equal sized end-chambers (20 cm x 18 cm x 35 cm) and a central chamber (20 cm x 10 cm x 35 cm). One end-chamber had a perforated floor and plain black walls, whereas the other end-chamber had a smooth floor and walls with vertical black and white stripes. The CPA protocol consisted of a pre-conditioning phase, four saline/morphine administration days in their home cages, a conditioning phase following morphine precipitated withdrawal in the CPA apparatus and a post-conditioning test. On Day 1 (pre-conditioning phase), mice were placed in the CPP apparatus and were allowed to explore all compartments for a period of 20 min. Naloxone-paired compartment was assigned to be in their most preferred compartment defined during the pre-conditioning test. On Days 2-5, mice were given a single injection of saline or morphine (5 mg/kg, s.c.) in their home cages (same time as their pre-conditioning test). During the conditioning session (i.e. Day 5, 1 hr and 50 min after their last saline or morphine injection), mice were given a single injection of saline or (*2R,6R*)-HNK (10 mg/kg, i.p.) and 10 min later were given another injection of either saline (4 ml/kg) or the opioid receptor antagonist naloxone (1 mg/kg, i.p.) and immediately placed in the CPA apparatus and confined in their preferred compartment (based on the pre-conditioning phase) for 30 min. Twenty-four hrs later, mice were placed in the CPA apparatus and were allowed to freely explore all three compartments for 20 min. Time spent in each compartment was measured during both pre- and post-conditioning sessions using TopScan v2.0 (CleverSys, Inc, VA, USA).

#### Experiment 4: Effect of (*2R,6R*)-HNK on precipitated morphine withdrawal somatic symptoms in opioid-dependent mice

Acute cessation of opioid use is characterized by short-lived somatic symptoms, including physical discomfort, diarrhea, abdominal pain, tremors, and a general reduced motivation to escape from aversive situations. Here, we investigated the potential therapeutic efficacy of (*2R,6R*)-HNK in mitigating acute morphine abstinence somatic withdrawal symptoms in mice.

### Precipitated morphine withdrawal symptoms

Mice were given a single injection of saline or morphine (5 mg/kg, s.c.) in their home cages for 4 days (one injection daily). On Day 5, 1 hr and 50 min after their last saline or morphine injection, mice were given a single injection of saline or (*2R,6R*)-HNK (10 mg/kg, i.p.) and 10 min later were given another injection of either saline or the opioid receptor antagonist naloxone (1 mg/kg, i.p.) and immediately placed in plexiglass cylinders and their behaviors were monitored and live-scored for 25 min by an experimenter blind to any treatment conditions. Behaviors that were scored included: number of jumps (escape behavior), number of paw tremors (1 point) or not in each 5-min bin for the following symptoms: stereotypic sniffing, stereotypic rearing, eye ptosis, piloerection and swallowing. An average score for each animal was given after analyzing all the withdrawal symptoms.

#### Experiment 5: Effect of (*2R,6R*)-HNK on maladaptive behaviors induced by protracted opioid withdrawal in mice

Although the somatic symptoms of opioid withdrawal are short-lived, individuals abstaining from opioid abuse suffer from emotional disturbances and other brain disorders, including depression, anxiety, and cognitive impairment, many years after detoxification. These behavioral disturbances act as a motivational trigger of relapse to opioid abuse. Here, we aimed to understand whether (*2R,6R*)-HNK is able to reverse maladaptive behavioral responses induced by protracted, 3-weeks, abstinence from chronic morphine administration in mice.

### Chronic escalating-dose morphine administration paradigm

Based on our previously published methods ^31^, mice were administered twice daily (10:00AM and 18:00PM) escalating doses of morphine (20, 40, 60, 80, 100 mg/kg) or saline (i.p.) for 5 days, followed by a single 100 mg/kg injection on day 6 (10:00AM). After this administration paradigm, mice were left drug-free for a period of 22 days (3 weeks; Day 28 of the protocol; Fig. xx). On that day, mice received a single injection of (*2R,6R*)-HNK (i.p.) or saline. On Day 29-30 (23-24 days withdrawal) mice were tested for sucrose preference overnight, female urine sniffing test (24 days withdrawal), social interaction (25 days withdrawal), anxiety-like behaviors in the Light/Dark box (26 days withdrawal) and spontaneous alternations in the Y-maze to measure spatial memory (27 days withdrawal).

### Sucrose preference (as previously published ^32^)

As described in Experiment 1.

### Female urine sniffing test (as previously published ^33^)

Mice were singly housed in freshly made home cages for a habituation period of 10 min. Subsequently, one plain cotton tip was secured on the center of the cage wall and mice were allowed to sniff and habituate to the tip for a period of 30 min. Then, the plain cotton tip was removed and replaced by two cotton tip applicators, one infused with fresh female mouse estrus urine and the other with fresh male mouse urine. These applicators were presented and secured at the two corners of the cage wall simultaneously. Sniffing time for both female and male urine was scored by a trained observer for a period of 2 min.

### Social interaction test

As described in Experiment 1.

### Light/dark box

Mice were placed in the illuminated compartment of the light/dark box (35 × 35 cm; Stoetling, USA), facing the wall opposite to the dark compartment. Mice were allowed to explore the whole apparatus for 10 min, as previously described ^34^. The sessions were recorded using overhead video cameras and the time spent in the illuminated and the dark compartments was scored for the first and last 5 min, by an experimenter blind to the experimental groups.

### Y-maze

Following measurement of sucrose and tap water consumed, mice were tested in the Y-maze for spatial memory testing. The procedure was performed based on our prior publications ^35^. Briefly, mice were placed into the “start” arm of the maze and thereafter explored the apparatus for 8 min. All the experimental trials were recorded with a digital video-camera. The videos were then scored by a trained observer blind to the experimental groups. Number of total arm entries and sequence of arm entries were measured. Percent alternations were calculated as: [the number of consecutive entries into three different compartments, divided by the total alternations (number of arm entries minus 2)] x 100.

#### Experiment 6: Effect of (*2R,6R*)-HNK on sub-threshold stress-induced behavioral impairment in mice previously exposed to morphine

A substantial body of evidence suggests that individuals with a history of opioid exposure are more vulnerable to stress-induced mood disturbances. To replicate and further investigate this phenomenon, we first aimed to determine whether prior exposure to a rewarding dose of morphine (5 mg/kg) can induce a susceptibility phenotype in mice undergoing a sub-threshold social defeat stress. We then sought to understand whether (*2R,6R*)-HNK prevents this behavioral impairment in mice that were exposed to opioids.

### Morphine administration

Mice were given a single daily injection of saline or morphine (5 mg/kg, s.c.) in their home cages, for a total of 4 days. Ten days after their last saline or morphine injection, mice underwent a subthreshold social defeat stress protocol. 2 hours after stress, mice were given a single injection of saline or (*2R,6R*)-HNK (10 mg/kg, i.p.).

### Sub-threshold social defeats

This experiment was performed as we previously published ^36^. On day 15, experimental mice were introduced to the home cage (43 cm length x 11 cm width x 20 cm height) of a resident aggressive retired CD-1 breeder - prescreened for aggressive behaviors - for 2 min. Following this physical attack phase, mice were transferred and housed in the opposite side of the resident’s cage divided by a perforated Plexiglas divider, in order to maintain sensory contact for 15 min. This process was repeated within the same day 3 times consecutively, with experimental mice being introduced to a novel aggressive CD-1 mouse for each of those 3 trials. Control mice (non-stressed) were placed in a new social defeat cage that did not contain a CD-1 aggressive mouse in a separate room.

### Social interaction test

The experiment was conducted as described in Experiment 5.

### Sucrose preference

The experiment was conducted as described in Experiment 5.

### Female urine sniffing test

After measuring sucrose (day 17), mice were placed (singly) in freshly made home cages for a habituation period of 10 min. The experiment was conducted as described in Experiment 5.

### Reinstatement of maladaptive behaviors using mild, sub-threshold stress exposure

Seven days after the female urine sniffing test (10 days after the sub-threshold social defeats) mice were re-screened for sucrose preference. All mice were then subjected to a 1-min exposure in the home-cage of an aggressive CD-1 mouse for assessing reinstatement of the previously acquired maladaptive behaviors (4 days after determination of sucrose preference restoration, i.e., Day 28). Following this exposure, mice were placed back in their home cages and presented with two identical bottles containing either tap water or 1% (w/v) sucrose solution for sucrose preference overnight. Next day, sucrose preference was determined again.

#### Experiment 7: Effect of (*2R,6R*)-HNK administration on the extinction progress from morphine conditioning

Extinction involves the gradual reduction and eventual elimination of learned drug-associated behaviors. Given the persistent challenges associated with opioid addiction, understanding how (*2R,6R*)-HNK may influence the speed of extinction from morphine conditioning is of significant interest.

### Morphine conditioning/extinction protocol

We used a morphine CPP protocol based on our previous studies ^37^. The protocol consisted of a habituation session, a pre-conditioning test, 4 conditioning sessions (morning: a 5 mg/kg, s.c., morphine injection and 5 hrs later a saline injection), a post-conditioning test and then once daily extinction sessions, with extinction progress testing in the morning of Days 2, 3 and 4 after the last morphine injection. We used the same CPP apparatus as described elsewhere in the manuscript. On Day 1, mice were placed in the CPP apparatus for a 20-min habituation period and were allowed to freely explore all chambers. We used a randomized, non-biased protocol. During the morning conditioning sessions, saline (4 ml/kg), or morphine (5 kg/kg) were administered, and mice were placed in on one of the two compartments (drug-paired compartment) to explore for 30 min. Five hours later mice were injected with saline (4 ml/kg) and were placed in the other compartment for 30 min. After the conditioning sessions, on Day 6, mice were tested for conditioning (post-Conditioning test). On Day 7, mice that developed conditioning to morphine were injected with (*2R,6R*)-HNK (10 mg/kg, i.p.) or 4 ml/kg saline, i.p. On Day 8, mice were injected with saline (4 ml/kg) in the morning and placed in their previously drug-paired compartment for 30 min and the same process was followed in the afternoon where mice were placed in their non-drug-paired compartment for 30 min. On Days 9, 10 and 11 (Day 2 Ext, Day 3 Ext and Day 4 Ext, respectively; Fig. 6A) mice were given free access to both compartments in the morning session (20 min total; to assess for Extinction progress) and were enclosed in their non-drug-paired compartment during the afternoon sessions after an injection of saline, to continue with the Extinction process. Sessions were videotaped via overhead digital cameras and were scored by a trained researcher blind to the treatment conditions.

#### Experiment 8: Effect of (*2R,6R*)-HNK administration on stress-induced reinstatement of morphine place preference

Here, we examined the efficacy of (*2R,6R*)-HNK on reversing stress-induced reinstatement of opioid-seeking behaviors following withdrawal.

### Morphine conditioned-place preference reinstatement protocol

We used the same Stoelting apparatus as described elsewhere in the manuscript. On Day 1, mice were placed in the CPP apparatus for a 20-min habituation period and were allowed to freely explore all chambers. We used a randomized, non-biased protocol. During the morning conditioning sessions, saline (4 ml/kg), or morphine (5 kg/kg) were administered s.c., and mice were placed in on one of the two compartments (drug-paired compartment) to explore for 30 min. Five hours later mice were injected s.c. with saline (4 ml/kg) and were placed in the other compartment for 30 min. After the conditioning sessions, on Day 6, mice were tested for conditioning (post-Conditioning test). On Days 7-12, mice were injected with saline (4 ml/kg) in the morning and placed in their previously drug-paired compartment for 30 min and the same process was followed in the afternoon where mice were injected with saline and placed in their non-drug-paired compartment for 30 min. On Day 13, mice were tested whether they had successfully extinct their morphine conditioning (post-Extinction test). Mice that successfully showed extinction were given an injection of (*2R,6R*)-HNK (10 mg/kg, i.p.) or 4 ml/kg saline, i.p. 3 hours after their extinction test session. On Day 14 (Reinstatement phase), mice were individually placed in a cylinder containing water (24±1 °C) and let to swim for 5 min for induction of stress, as we previously published ^31^. After exposure to the swim stress, mice were thoroughly dried using paper towels and placed in the CPP chambers to explore both compartments, as we have described previously ^31^. Sessions were videotaped via overhead digital cameras and were scored by a researcher blind to the treatment conditions.

#### Experiment 9: Effect of (*2R,6R*)-HNK administration on voluntary consumption of morphine in mice that were exposed to non-contingent administration of opioids

Here, we established a novel mouse model designed to replicate the human behavioral response, wherein prior exposure to opioids, possibly through prescriptions, leads to an increased tendency toward opioid use upon subsequent exposure. More specifically, we exposed mice to our non-contingent escalating morphine administration paradigm (described in Experiment 5). Forty-eight hours after the last morphine injection mice received a saline or (*2R,6R*)-HNK (10 mg/kg) injection let mice withdraw from morphine for a period of 3 weeks. After that (Day 29 of the protocol), mice were presented with two identical bottles containing either tap water + 0.1 mg/ml quinine or tap water + 0.1 mg/ml quinine + 0.3 mg/ml morphine. This 2-bottle choice morphine protocol was adapted from Grim et al., (2019)^38^. To ensure that morphine preference is not due to taste differences, after 4 days of testing for oral morphine preference, we removed the morphine choice and gave 2 identical 0.1 mg/ml quinine bottles (Day 5). Morphine and quinine bottles were weighed daily in the morning and positions of the bottles in the cages were interchanged daily to avoid place conditioning. Percent morphine preference was calculated as the amount of morphine liquid consumed divided by the total liquid amount consumed (morphine and quinine solutions) x 100. Morphine consumption was calculated by multiplying the 0.3 mg of morphine by the morphine liquid consumed per day and then dividing by the weight of each animal (in g) x 1000 to get mg/kg.

#### Experiment 10: Effect of (*2R,6R*)-HNK administration on markers of synaptic plasticity during protracted opioid withdrawal

### Western blot protocol

To assess the involvement of markers of synaptic plasticity in the effects of HNK to reverse protracted opioid-withdrawal induced maladaptive behavioral responses in mice, mice have undergone our 6-day escalating-dose morphine administration paradigm and protracted, 28-day withdrawal period (as described in Experiment 5). On Day 28 of withdrawal, mice received a single injection of saline or (*2R,6R*)-HNK (10 mg/kg). On day 29 of withdrawal, mice were euthanized via decapitation and brains were excised, and ventral hippocampi immediately dissected and preserved in -80°C until further analysis.

To purify the synaptoneurosomes, mouse ventral hippocampi were dissected and homogenized in Syn-PER Reagent (ThermoFisher Scientific; #87793) with 1X protease and phosphatase inhibitor cocktail (ThermoFisher Scientific; #78440). The homogenate was centrifuged for 10 min at 1,200 x g at 4°C. The supernatant was then centrifuged at 15,000 x g for 20 min at 4°C. After centrifugation, the pellet (synaptosomal fraction) was re-suspended in N-PER Neuronal Protein Extraction Reagent (ThermoFisher Scientific; 87792). The homogenate was transferred to an appropriate microcentrifuge tube and incubated on ice for 10 minutes. The sample was centrifuged at 10,000 × g for 10 minutes at 4°C and the supernatant was collected. Protein concentration was determined via Bradford assay (Merck; #1.10306). Equal amounts of proteins (20–100μg as optimal for each antibody) for each sample was loaded into 4-15% Mini-PROTEAN TGX Stain-Free Protein Gels (BioRad; #4568084) for electrophoresis. Gel transfer was performed with the Mini Trans-Blot® Transfer System (Bio-Rad; 1703930) Supported Nitrocellulose membranes (BioRad; #1620094). Transferred proteins were blocked with 5% milk in TBST (TBS plus 0.1% Tween-20) for 1h and incubated in primary antibodies (1:1000 in 5% BSA) overnight at 4°C. The following primary antibodies were use: phospho-eEF2 (at Thr56; Cell Signalling Technology; #2331), total eEF2 (Cell Signalling Technology; #2343), phospho-mTOR (at Ser2448; Cell Signalling Technology; #2971), total mTOR (Cell Signalling Technology; #2983), phospho-GluA1 (at Ser831; Cell Signalling Technology; #75574), GluA1 (Cell Signalling Technology; #13185), phospho-GluN2A (at Tyr1246; Cell Signalling Technology; #4206), GluN2A (Cell Signalling Technology; #4205), GluN1 (Cell Signalling Technology; #5784), BDNF (Cell Signalling Technology; #47808), and GAPDH (Cell Signalling Technology; #97166). The next day, blots were washed three times in TBST and incubated with horseradish peroxidase conjugated anti-mouse or anti-rabbit secondary antibody (1:1000 to 1:3000 in 5% milk) for 1h. After three final washes with TBST, bands were detected using enhanced chemiluminescence (ECL) (BioRad; 1705060) with the ChemiDoc XRS+ System (BioRad). After imaging, the blots were incubated in stripping buffer (ThermoFisher Scientific; 46430) for 15 min at room temperature followed by three washes with TBST. The stripped blots were incubated in blocking solution (5% milk) for 1h and incubated with the primary antibody directed against total levels of the respective protein or GAPDH for loading control. Densitometric analysis of phospho- and total immunoreactive bands for each protein was conducted using ImageJ (Fiji) software. The values for the phosphorylated forms of proteins were normalized to phosphorylation-independent levels of the same protein. Phosphorylation-independent levels of proteins were normalized to GAPDH. Fold change was calculated by normalization to saline-treated control group for each protein or phosphoprotein.

### Statistical Analyses

Required samples sizes were estimated based upon our past experience performing similar experiments. Experimentation and analysis were performed in a manner blind to treatment assignments in all experiments with the exception of the whole-cell patch-clamp recordings and intravenous drug self-administration. For all blinded experiments, mice were randomly assigned to treatment groups. Statistical analyses were performed using GraphPad Prism software v9. All statistical tests were two-tailed, and significance was assigned at p < 0.05. Normality and equal variances between group samples were assessed using the Kolmogorov-Smirnov and Brown-Forsythe tests respectively. ANOVAs were followed by a Holm-Šídák *post-hoc* comparison when significance was reached, and significant results are indicated with asterisks in the Figures.

## RESULTS

### Experiment 1: Administration of (*2R,6R*)-HNK prevents the development of conditioning to sub-effective doses of morphine in stress susceptible mice

Stress susceptible mice, determined by <50% social interaction preference and <55% sucrose preference, (but not resilient mice) developed place preference to a sub-effective dose of morphine (i.e., 0.25 mg/kg), indicating that stress susceptibility for developing anhedonia and sociability deficits enhances the probability of developing conditioning to opioids at very low, sub-effective doses (**Fig. 1**). Notably, a single administration of (*2R,6R*)-HNK 24 hrs prior to the initiation of the conditioning phase of the CPP, prevented low dose morphine-induced place preference in stress susceptible mice, with no significant effect in stress-resilient mice [Resilient mice: 2-way RM ANOVA, Experimental Group effect: F (3, 40) = 0,3001, P=0,8251; CPP phase effect: F (1, 40) = 1,984, P=0,1667; Interaction effect: F (3, 40) = 0,6104, P=0,6122; n= 11/group; Susceptible mice: 2-way RM ANOVA, Experimental Group effect: F (3, 37) = 0,8817, P=0,4595; CPP phase effect: F (1, 37) = 6,466, P=0,0153; Interaction effect: F (3, 37) = 2,720, P=0,0583; n= 10-11/group] (**Fig. 1**).

**Figure 1.**
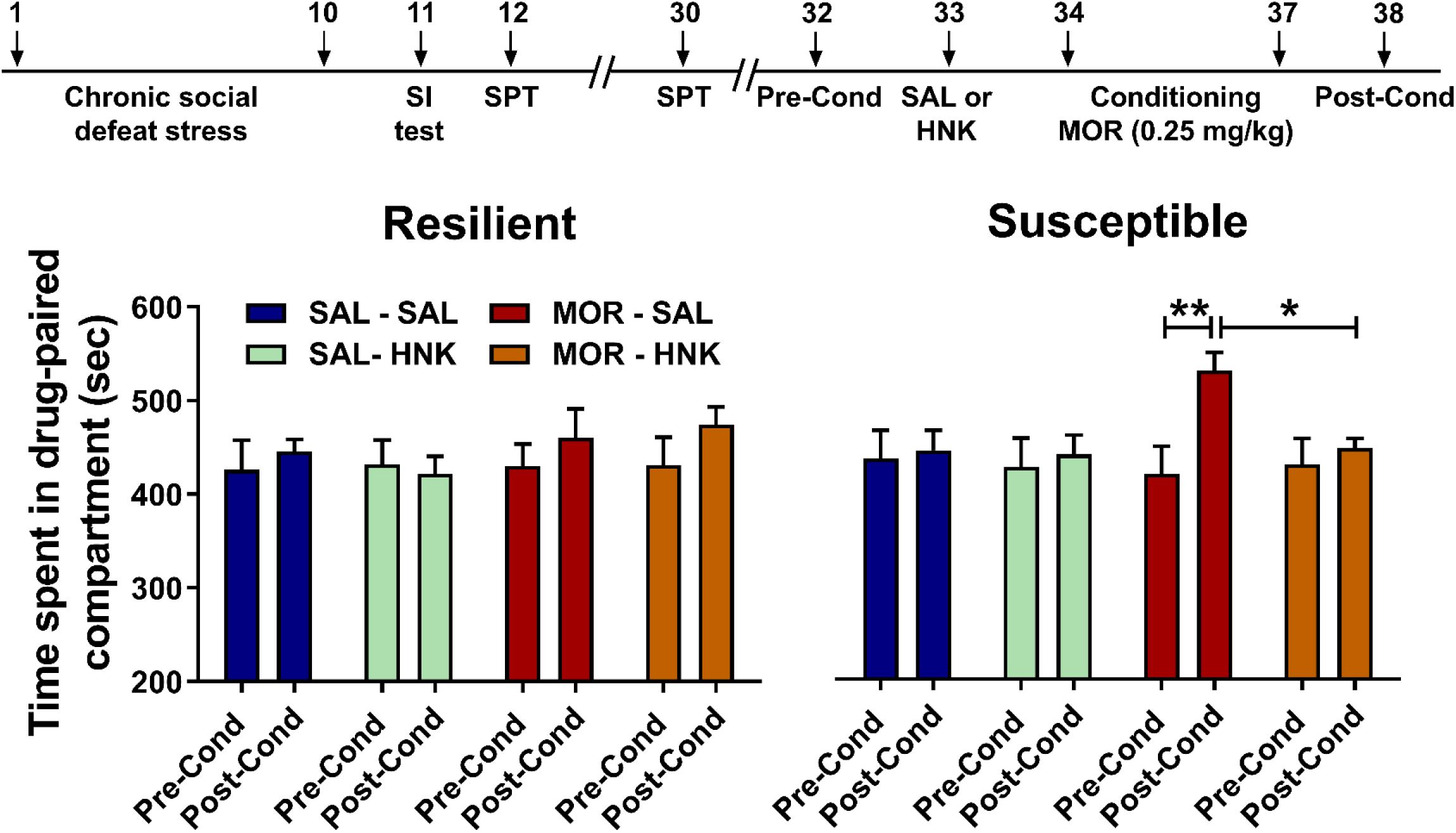
Administration of (*2R,6R*)-HNK prevents the development of conditioning to sub-effective doses of morphine in stress susceptible mice. Stress-susceptible mice, characterized by <50% social interaction preference and <55% sucrose preference, developed a place preference to a low, sub-effective dose of morphine (0.25 mg/kg). This suggests that stress susceptibility, linked to anhedonia and sociability deficits, heightens the likelihood of conditioning to opioids even at very low doses. Notably, a single administration of (*2R,6R*)-HNK (10 mg/kg) twenty-four hours before initiating the conditioning phase prevented this low-dose morphine-induced place preference exclusively in stress-susceptible mice. Data are the mean ± S.E.M.; n= 10-11 mice/group; * *p*<0.05, ** *p*<0.01. *<u>Abbreviations</u>*: Cond, Conditioning; HNK, hydroxynorketamine; MOR, morphine; SAL, saline; SPT, sucrose preference test; SI test, social interaction test.

### Experiment 2: Administration of (*2R,6R*)-HNK does not prevent the development of conditioning to high, rewarding doses of morphine

A single dosing of (*2R,6R*)-HNK did not prevent the development of high-dose (5 mg/kg) morphine CPP in mice [,2-way RM ANOVA, Experimental Group effect: F (3, 28) = 0,9547, P=0,4277; CPP phase effect: F (1, 28) = 14,12, P=0,0008; Interaction effect: F (3, 28) = 6,032, P=0,0027; n= 8/group] (**Supplementary Fig. 1**), indicating that in Experiment 1, (*2R,6R*)**-**HNK possibly reversed the stress-susceptibility phenotype of mice with a secondary effect to reverse conditioning to sub-effective doses of morphine and not morphine conditioning *per se*.

### Experiment 3: Administration of (*2R,6R*)-HNK prevents conditioned-place aversion induced by precipitated morphine withdrawal in opioid-dependent mice

Administration of (*2R,6R*)-HNK prevented the development of naloxone-induced place aversion in morphine-dependent mice [2-way ANOVA, Treatment effect (SAL vs MOR): F (1, 72) = 0,01253, P=0,9112; Experimental Groups effect: F (3, 72) = 5,759, P=0,0014; Interaction effect: F (3, 72) = 2,056, P=0,1137; n= 10/group] (**Fig. 2**).

**Figure 2.**
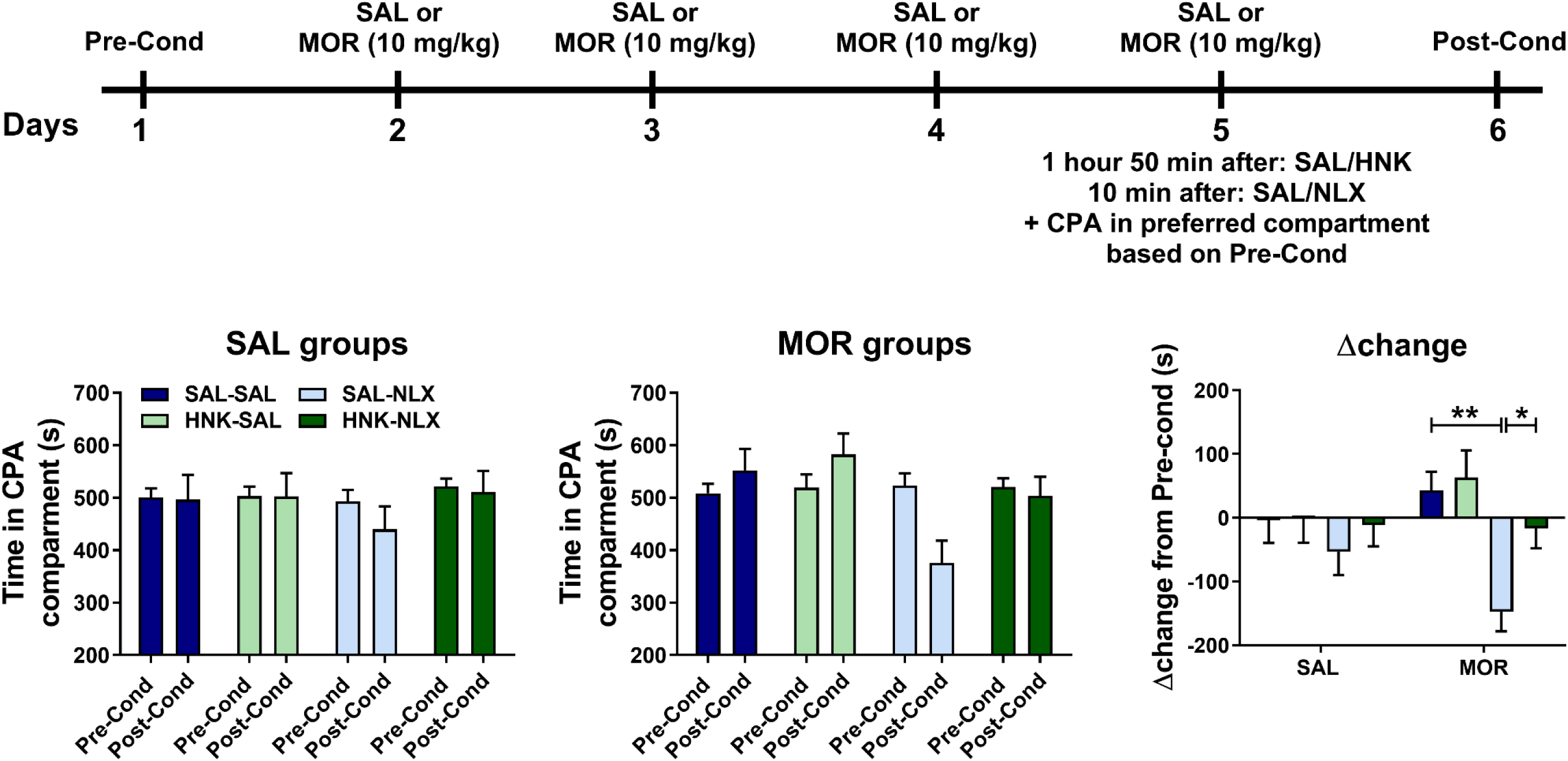
Administration of (2R,6R)-HNK prevents conditioned-place aversion induced by precipitated morphine withdrawal in opioid-dependent mice. To evaluate the therapeutic potential of (2R,6R)-HNK in mitigating negative affective states linked to acute opioid abstinence, mice were rendered dependent on morphine and subsequently administered the opioid receptor antagonist naloxone to induce precipitated withdrawal. Notably, (*2R,6R*)-HNK (10 mg/kg) administration completely blocked the development of naloxone-induced place aversion in morphine-dependent mice. Data are the mean ± S.E.M.; n= 9 mice/group; * *p*<0.05, ** *p*<0.01. *<u>Abbreviations</u>*: Cond, Conditioning; CPA, conditioned-place aversion; HNK, hydroxynorketamine; MOR, morphine; NLX, naloxone; SAL, saline.

### Experiment 4: Administration of (*2R,6R*)-HNK enhances escape behavior and reduces overall precipitated morphine withdrawal symptoms in opioid-dependent mice

Administration of (*2R,6R*)-HNK while increasing the escape-related jumping behavior [2-way ANOVA: Main treatment effect: F (1, 103) = 82,58, P<0,0001; HNK treatment effect: F (3, 103) = 33,19, P<0,0001; Interaction effect: F (3, 103) = 33,11, P<0,0001], it decreased paw tremors [2-way ANOVA: Main treatment effect: F (1, 101) = 74,77; P<0,0001; HNK treatment effect: F (3, 101) = 45,25; P<0,0001; Interaction effect: F (3, 101) = 29,41, P<0,0001] and eye ptosis [2-way ANOVA: Main treatment effect: F (1, 101) = 57,85; P<0,0001; HNK treatment effect: F (3, 101) = 36,44, P<0,0001; Interaction effect: F (3, 101) = 27,93, P<0,0001] and induced an overall reduction in average precipitated withdrawal symptoms of morphine [2-way ANOVA: Main treatment effect: F (1, 104) = 127,0, P<0,0001; HNK treatment effect: F (3, 104) = 78,26, P<0,0001; P<0,0001; Interaction effect: F (3, 104) = 47,86, P<0,0001] (**Fig. 3**). No significant effect was observed in any other somatic symptoms scored. For all measures there were 13-14 mice per experimental group.

**Figure 3.**
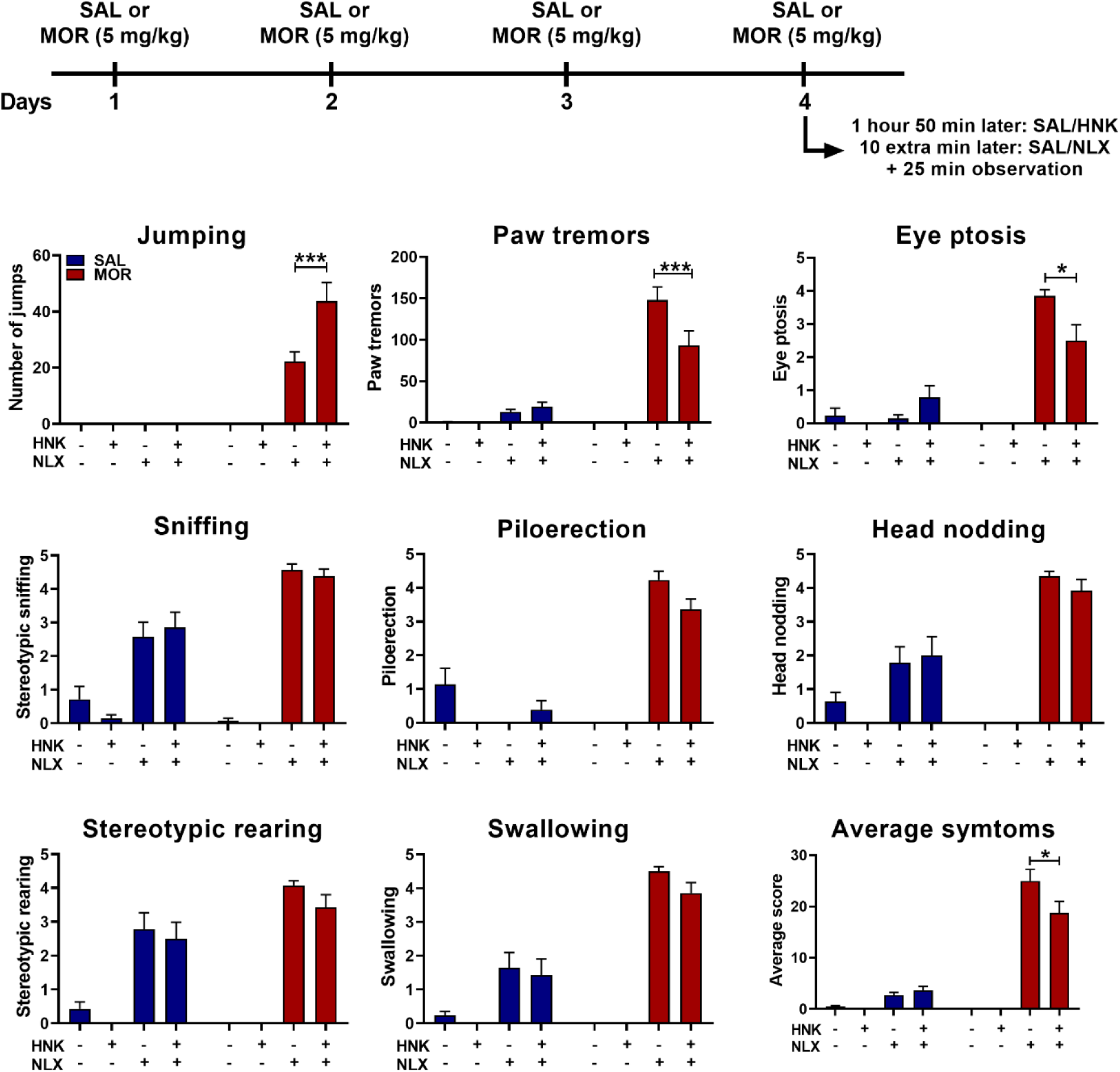
Administration of (*2R,6R*)-HNK reduces the somatic symptoms induced by acute, precipitated opioid withdrawal. (*2R,6R*)-HNK administration demonstrated significant improvements in escape behavior, reducing paw tremors, eye ptosis, and overall morphine withdrawal symptoms induced by administration of naloxone (NLX; 1 mg/kg) in opioid-dependent mice. Importantly, (*2R,6R*)-HNK notably increased escape-related jumping behavior. Data are the mean ± S.E.M.; n=13-14 mice/group; * *p*<0.05, *** *p*<0.001. *<u>Abbreviations</u>*: HNK, hydroxynorketamine; MOR, morphine; NLX, naloxone; SAL, saline.

### Experiment 5: Single administration of (*2R,6R*)-HNK reverses maladaptive behaviors induced by protracted opioid withdrawal in mice

A single administration of (*2R,6R*)-HNK reversed anhedonia phenotypes, as indicated by normalization of protracted abstinence-indued reductions in sucrose preference [2-way ANOVA: Treatment effect: F (1, 28) = 9,916, P=0,0039; HNK effect: F (1, 28) = 4,789, P=0,0371; Interaction effect: F (1, 28) = 11,73, P=0,0019; n=8/group] (**Fig. 4B**), female urine sniffing preference [2-way ANOVA: Treatment effect: F (1, 28) = 7,466, P=0,0108; HNK effect: F (1, 28) = 9,437, P=0,0047; Interaction effect: F (1, 28) = 16,94; P=0,0003; n=8/group] (**Fig. 4C**), social interaction deficits [2-way ANOVA: Treatment effect: F (1, 28) = 2,325, P=0,1386; HNK effect: F (1, 28) = 3,769, P=0,0623; Interaction effect: F (1, 28) = 5,706, P=0,0239; n=8/group] (**Fig. 4D**), anxiety-like behaviors in the light/dark box [2-way RM ANOVA: Treatment effect: F (3, 26) = 4,921, P=0,0077; Time in the L/D box effect: F (1, 26) = 30,45, P<0,0001; Interaction effect: F (3, 26) = 1,707, P=0,1902; n=8/group] (**Fig. 4E**) and cognitive impairment in the Y-maze [2-way ANOVA: Treatment effect: F (1, 28) = 5,158, P=0,0310; HNK effect: F (1, 28) = 7,535, P=0,0104; Interaction effect: F (1, 28) = 11,70, P=0,0019; n=8/group] (**Fig. 4F**), induced by protracted, 3-week, morphine withdrawal.

**Figure 4.**
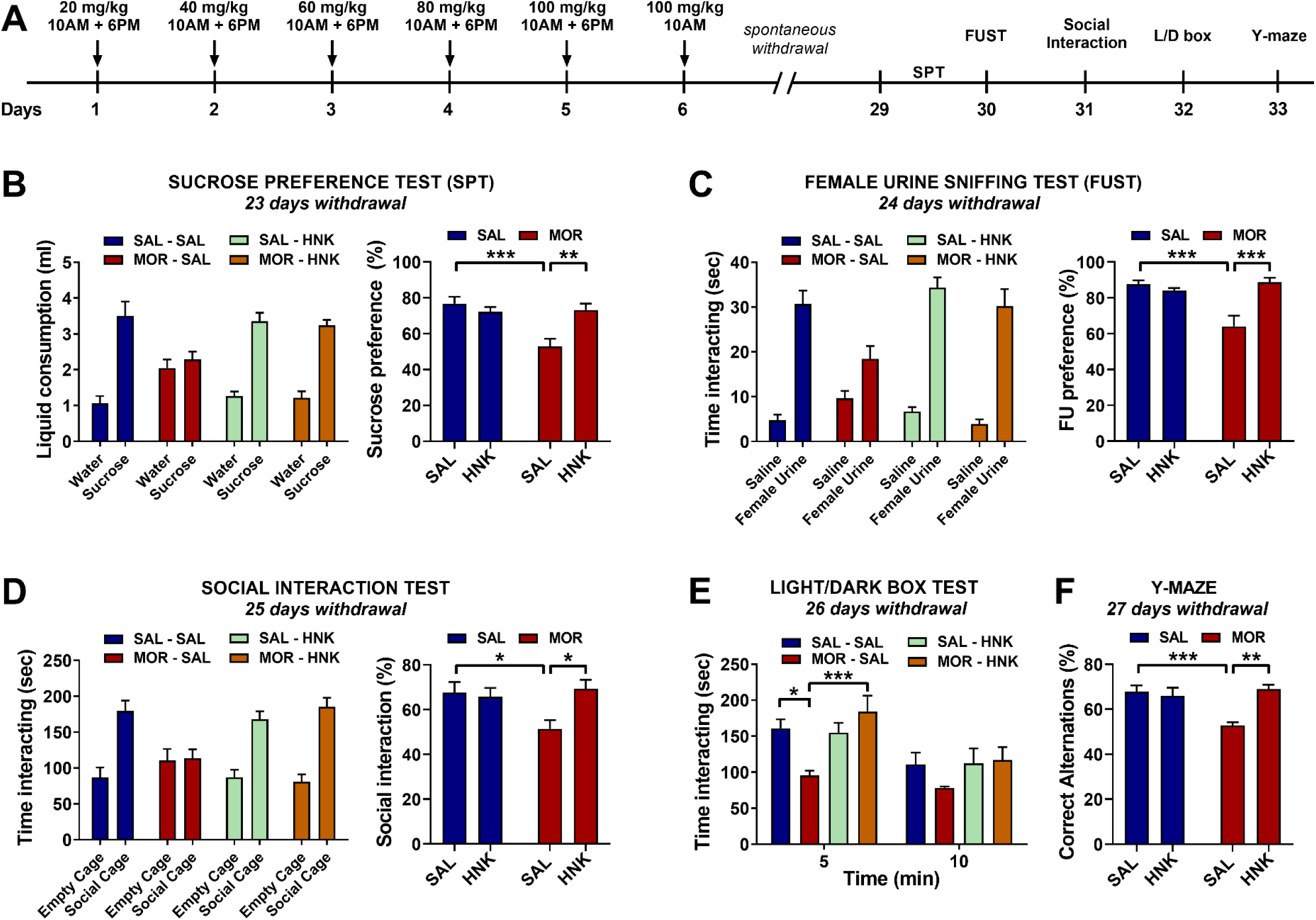
Administration of (*2R,6R*)-HNK prevents maladaptive behavioral responses induced by protracted morphine withdrawal in mice. **(A)** Timeline for the experimental procedures. (*2R,6R*)-HNK administration effectively restored reduced **(B)** sucrose preference, **(C)** female urine sniffing preference, **(D)** social interaction deficits, and **(E)** anxiety-like behaviors in the light/dark box, and **(F)** improved cognitive impairment induced by a 3-week morphine withdrawal. Data are the mean ± S.E.M.; n= 8 mice/group; * *p*<0.05, ** *p*<0.01, *** *p*<0.001. *<u>Abbreviations</u>*: HNK, hydroxynorketamine; L/D box, Light/Dark box; MOR, morphine; SAL, saline.

### Experiment 6: Single administration of (*2R,6R*)-HNK reverses stress susceptibility for developing maladaptive behaviors induced by previous exposure to opioids and protects from a subsequent stress re-exposure

A single administration of (*2R,6R*)-HNK reversed stress susceptibility phenotypes of mice previously exposed to morphine in the social interaction [2-way ANOVA: Treatment effect: F (1, 71) = 2,922, P=0,0917; HNK effect: F (3, 71) = 6,710, P=0,0005; Interaction effect: F (3, 71) = 4,922, P=0,0037; n=9-10/group], sucrose preference [2-way ANOVA: F (1, 71) = 6,715, P=0,0116; HNK effect: F (3, 71) = 3,663, P=0,0163; Interaction effect: F (3, 71) = 2,397, P=0,0752; n=9-10/group], female urine sniffing [2-way ANOVA: F (1, 71) = 2,636, P=0,1089; HNK effect: F (3, 71) = 4,950, P=0,0035; Interaction effect: F (3, 71) = 3,534, P=0,0190; n=9-10/group] and nestlet building tests [2-way ANOVA: F (1, 71) = 4,490, P=0,0376; HNK effect: F (3, 71) = 4,480, P=0,0062; Interaction effect: F (3, 71) = 3,018, P=0,0354; n=9-10/group] (**Fig. 5A**).

**Figure 5.**
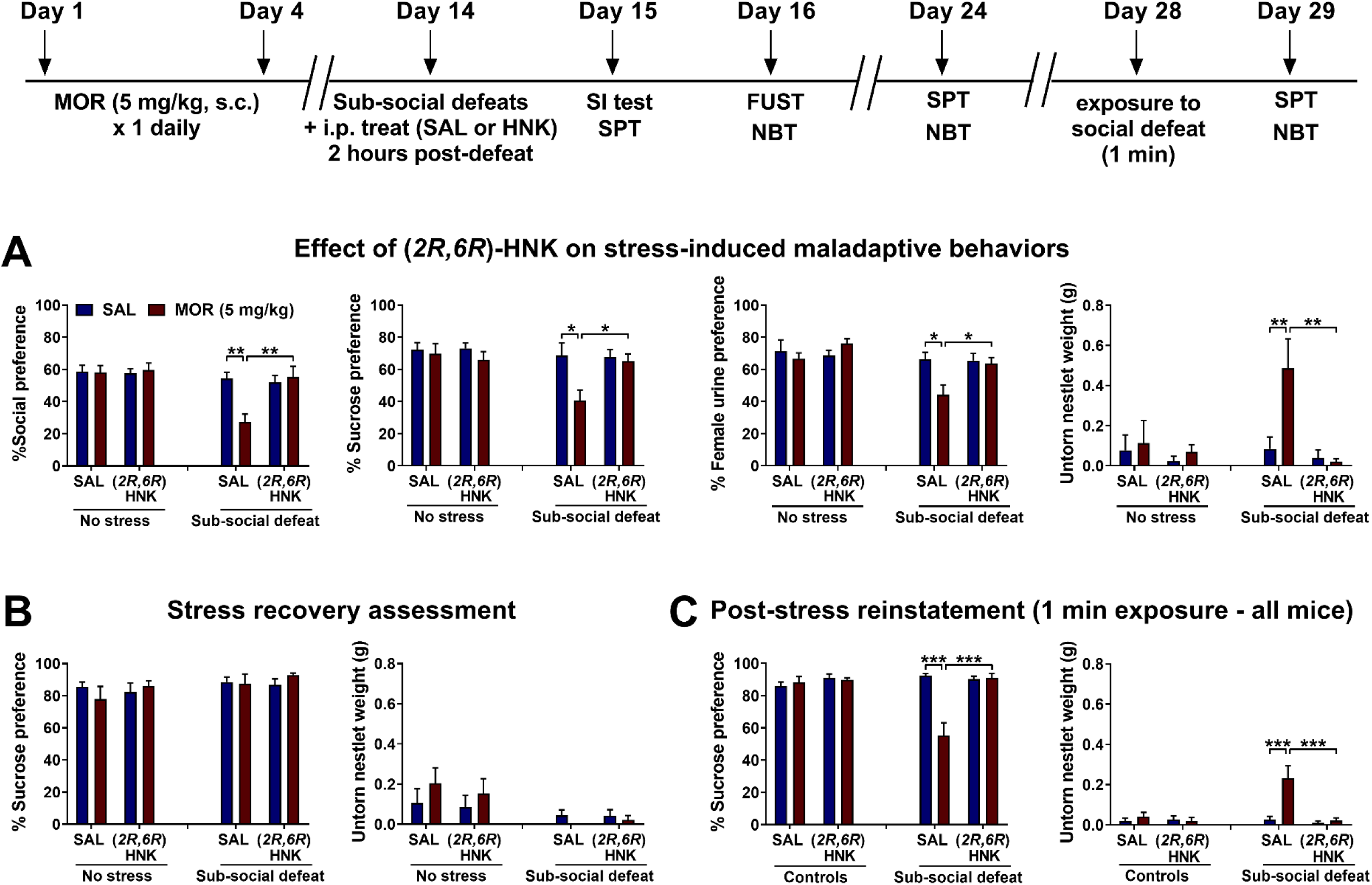
Administration of (*2R,6R*)-HNK to mice previously exposed to opioids prevented maladaptive behavioral responses induced by exposure to a sub-threshold social defeat stress. Prior exposure to a rewarding dose of morphine (5 mg/kg, s.c.) rendered mice susceptible to a subsequent sub-threshold social defeat stress-induced behavioral impairment. **(A)** (*2R,6R*)-HNK administration (10 mg/kg) reversed stress susceptibility phenotypes in these mice, restoring social interaction, sucrose preference, female urine sniffing, and nestlet building behaviors. **(B)** A subsequent 8-day interval, spontaneously reversed anhedonia and reduced nestlet building behaviors induced by prior opioid exposure and stress. **(C)** Subsequent mild stress exposure reinstated maladaptive behaviors in saline-treated animals but not in (*2R,6R*)-HNK-treated mice. Data are the mean ± S.E.M.; n= 9-10 mice/group; * *p*<0.05, ** *p*<0.01, *** *p*<0.001. *<u>Abbreviations</u>*: FUST, female urine sniffing test; HNK, hydroxynorketamine; MOR, morphine; NBT, nestlet building test; SAL, saline; SI test, social interaction test; SPT, sucrose preference test.

**Figure 6.**
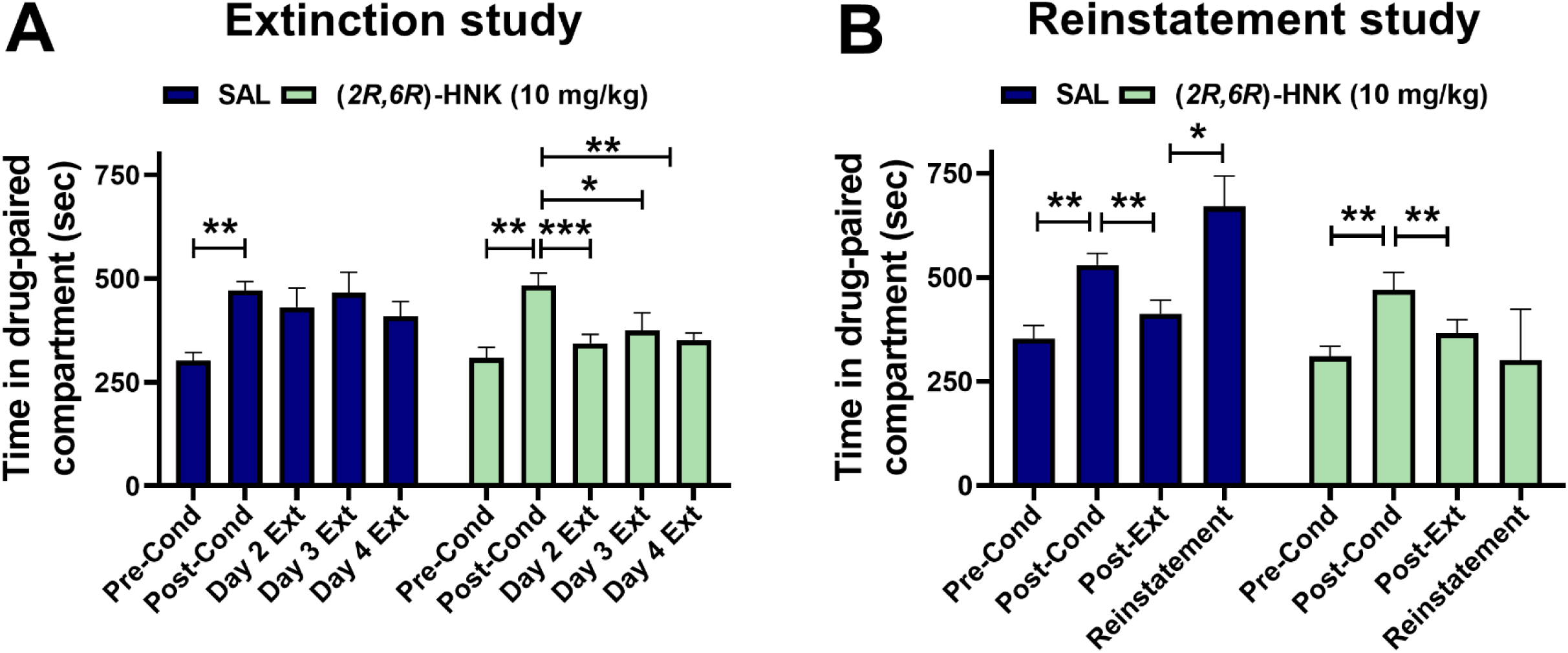
Administration of (*2R,6R*)-HNK facilitates extinction from opioid conditioning and prevents stress-induced reinstatement in mice. **(A)** (*2R,6R*)-HNK administration (10 mg/kg) induced complete extinction of morphine conditioning two days after initiation of the extinction period in the conditioned-place preference paradigm, contrasting with saline-treated mice, which did not achieve complete extinction even four days post-abstinence. **(B)** Additionally, in a different cohort of mice (*2R,6R*)-HNK administration prevented stress-induced reinstatement of morphine conditioning behavior after seven days of abstinence. Data are the mean ± S.E.M.; n= 7-9 mice/group; * *p*<0.05, ** *p*<0.01, *** *p*<0.001. *<u>Abbreviations</u>*: Cond, conditioning; Ext, extinction; HNK, hydroxynorketamine; SAL, saline.

After 8 days mice were re-tested in the sucrose preference test and nestlet building behaviors. Results showed that this time was adequate to reverse anhedonia phenotypes induced by a double hit paradigm throughprior exposure to opioids and a subthreshold stressor (P>0.05; **Fig. 5B**).

Following an additional 4 days, all mice were exposed to a very mild, transient (1 min) exposure to social defeat attack to assess for possible prophylactic effects of (*2R,6R*)-HNK. Indeed, 1 min stress exposure reinstated maladaptive behaviors in saline-treated animals, but not (*2R,6R*)-HNK-treated mice [sucrose preference: 2-way ANOVA: F (1, 71) = 11,77, P=0,0010; HNK effect: F (3, 71) = 9,228, P<0,0001; Interaction effect: F (3, 71) = 13,20, P<0,0001; n=9-10/group; nestlet building behavior: 2-way ANOVA: F (1, 71) = 9,117, P=0,0035; HNK effect: F (3, 71) = 7,583, P=0,0002; Interaction effect: F (3, 71) = 6,681, P=0,0005; n=9-10/group] (**Fig. 5C**).

### Experiment 7: Single administration of (*2R,6R*)-HNK facilitates extinction progress from opioid conditioning

A single administration of (*2R,6R*)-HNK 8 hours following the last morphine injection facilitated extinction of morphine conditioning, with a total extinction occuring 2 days following morphine cessation, while saline-treated mice did not reach complete extinction even at 4 days after abstinence [2-way RM ANOVA: Treatment effect: F (1, 15) = 1,656, P=0,2177; CPP phase effect: F (2,918, 43,77) = 12,71, P<0,0001; Interaction effect: F (4, 60) = 3,032, P=0,0614; n=9-10/group; HNK effect: F (3, 71) = 7,583, P=0,0002; Interaction effect: F (3, 71) = 6,681, P=0,0005; n=8-9/group] (**Fig. 6A**).

### Experiment 8: Single administration of (*2R,6R*)-HNK prevents stress-induced reinstatement to morphine conditioning behavior after abstinence

A single administration of (*2R,6R*)-HNK 8 hrs following the last post-extinction test (7 days after final morphine injection), prevented swim stress-induced reinstatement of morphine conditioning behavior [2-way RM ANOVA: Treatment effect: F (1, 12) = 6,507, P=0,0254; CPP phase effect: F (1,218, 14,62) = 4,743, P=0,0402; Interaction effect: F (4, 60) = 3,032, P=0,0614; n=9-10/group] (**Fig. 6B**).

### Experiment 9: (*2R,6R*)-HNK prevents enhanced preference for opioids in mice previously exposed to non-contingent morphine administration

A single administration of (*2R,6R*)-HNK 48 hrs following the last morphine injection was able to decrease voluntary morphine preference [2-way RM ANOVA: Treatment effect: F (3, 36) = 5,109, P=0,0048; Time effect: F (3,422, 123,2) = 10,94, P<0,0001; Interaction effect: F (12, 144) = 0,7415, P=0,7090; n=10/group] (**Fig. 7A**) and general morphine consumption [2-way RM ANOVA: Treatment effect: F (3, 36) = 4,521, P=0,0086; Time effect: F (2,650, 95,40) = 7,940, P=0,0002; Interaction effect: F (9, 108) = 0,8468, P=0,5749; n=10/group] (**Fig. 7B**) in mice that were previously exposed to forced use of opioids.

**Figure 7.**
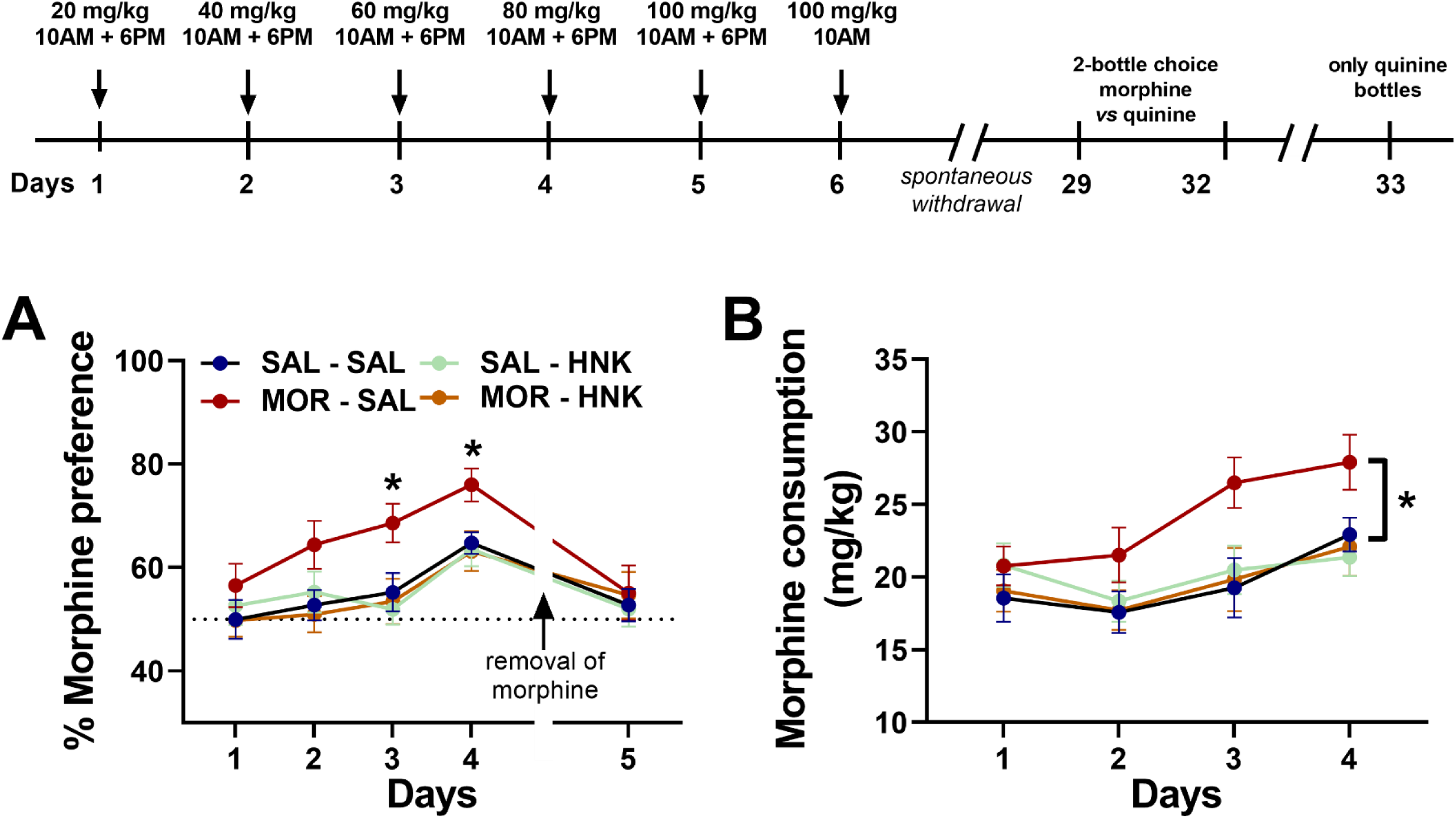
Administration of (2R,6R)-HNK prevents heightened preference for opioids in mice previously exposed to non-contingent morphine. Mice, subjected to a non-contingent escalating morphine administration paradigm, exhibited increased preference for morphine consumption after withdrawal. (*2R,6R*)-HNK (10 mg/kg) administered 48 hours post-morphine exposure significantly decreased voluntary **(A)** morphine preference and **(B)** overall morphine consumption. This suggests the potential of (2R,6R)-HNK in reducing heightened opioid preference induced by prior forced opioid use. Data are the mean ± S.E.M.; n= 10 mice/group; * *p*<0.05, ** *p*<0.01, *** *p*<0.001. *<u>Abbreviations</u>*: Cond, conditioning; Ext, extinction; HNK, hydroxynorketamine; MOR, morphine; SAL, saline.

### Experiment 10: Administration of (*2R,6R*)-HNK reverses reductions in BDNF and GluN2A/NMDARs measured in synaptoneurosomal ventral hippocampal fractions

Protracted withdrawal from morphine or administration of (*2R,6R*)-HNK did not alter GluA1 or phospho-GluA1 levels (P>0.05; **Fig. 8A**). In contrast, protracted morphine abstinence induced a significant decrease in the levels of GluN2A subunits of the NMDARs (but not phospho-GluN2A), while administration of (*2R,6R*)-HNK reversed this reduction [2-way ANOVA: Main treatment effect: F (1, 28) = 12,79, P=0,0013; HNK effect: F (1, 28) = 0,3931, P=0,5358; Interaction effect: F (1, 28) = 9,708, P=0,0042; n=8/group] (**Fig. 8B**). Protracted withdrawal from morphine or administration of (*2R,6R*)-HNK did not alter eEF2 or phospho-eEF2 levels (P>0.05; **Fig. 8A**). Similarly, protracted withdrawal from morphine or (*2R,6R*)-HNK did not alter mTOR levels (P>0.05; **Fig. 8D**), but morphine withdrawal induced a decrease in the phospho-mTOR [2-way ANOVA: Main treatment effect: F (1, 28) = 5,111, P=0,0317; HNK effect: F (1, 28) = 1,083, P=0,3068; Interaction effect: F (1, 28) = 0,2885, P=0,5954; n=8/group] (**Fig. 8D**). Morphine abstinence also decreased GluN1 [2-way ANOVA: Main treatment effect: F (1, 28) = 3,872, P=0,0591; HNK effect: F (1, 28) = 0,7864, P=0,3827; Interaction effect: F (1, 28) = 1,342, P=0,2564; n=8/group] levels, however, (*2R,6R*)-HNK did not reverse this effect (**Fig. 8E**). Finally, protracted morphine abstinence induced a significant decrease in the levels of BDNF, an effect that (*2R,6R*)-HNK reversed [2-way ANOVA: Main treatment effect: F (1, 28) = 3,680, P=0,0653; HNK effect: F (1, 28) = 0,8132, P=0,3749; Interaction effect: F (1, 28) = 4,306, P=0,0473; n=8/group] (**Fig. 8F**).

**Figure 8.**
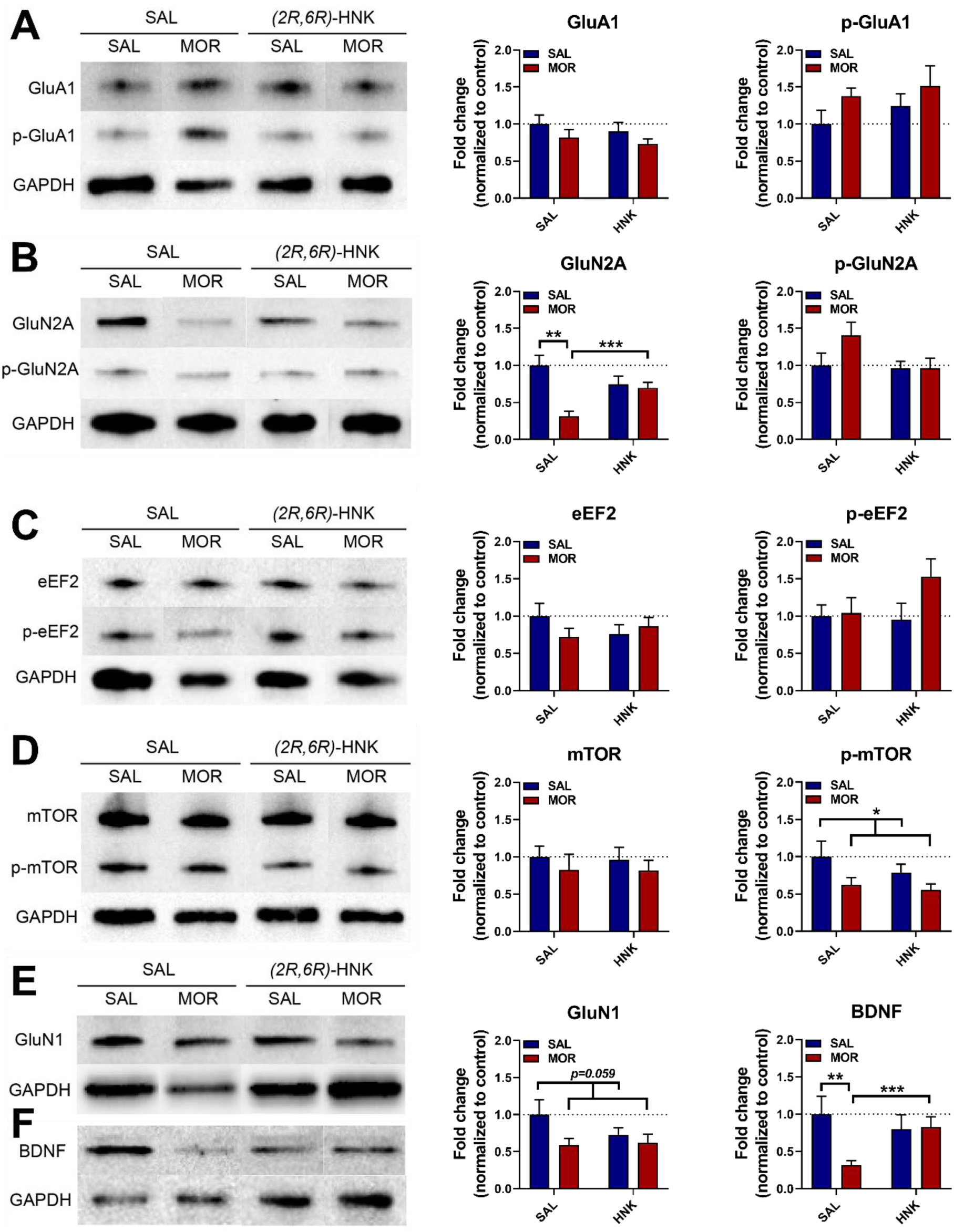
Administration of (*2R,6R*)-HNK reverses impaired synaptic plasticity markers in the ventral striatum. Following a 28-day withdrawal period from morphine, mice received a single injection of saline or (*2R,6R*)-HNK (10 mg/kg). Subsequent western blot analysis of synaptoneurosomal fractions of ventral hippocampus of these mice revealed that protracted morphine abstinence induces significant reductions in GluN2A subunits of the NMDARs and brain-derived neurotrophic factor (BDNF) levels, both of which (*2R,6R*)-HNK reversed. However, morphine withdrawal-induced alterations in phospho-mTOR and GluN1 were not reversed by (*2R,6R*)-HNK. No significant effects were observed in other measured markers of synaptic plasticity. Data are the mean ± S.E.M.; n= 10 mice/group; * *p*<0.05, ** *p*<0.01, *** *p*<0.001. *<u>Abbreviations</u>*: BDNF, brain-derived neurotrophic factor; eEF2, eukaryotic elongation factor 2; GluA1, GluA1 AMPA receptor subunit; GluN1, GluN1 NMDA receptor subunit; GluN2A, GluN2A NMDA receptor subunit; HNK, hydroxynorketamine; MOR, morphine; mTOR, mechanistic target of rapamycin; SAL, saline.

## DISCUSSION

In the present study, we developed innovative mouse models mimicking human behaviors associated with opioid addiction, specifically targeting stress susceptibility and conditioned responses to varying doses of morphine. These models allowed us to explore the interplay between stress vulnerability and opioid conditioning, offering insights into the complex dynamics of addiction development and potential treatment interventions. Across a series of experiments utilizing these models, we investigated the effects of (*2R,6R*)-HNK, a metabolite of ketamine, on various facets of opioid addiction and associated behaviors. We showed that (*2R,6R*)-HNK’s administration prevents conditioning to sub-effective doses of morphine in stress-susceptible mice, indicating its efficacy in mitigating opioid conditioning in stress vulnerable phenotypes. The fact that it did not prevent conditioning to higher morphine doses emphasizes its possible role in reversing stress susceptibility rather than directly altering morphine conditioning properties itself. Notably, (*2R,6R*)-HNK completely prevented naloxone-induced conditioned place aversion in morphine-dependent mice, and ameliorated precipitated withdrawal-related somatic symptoms, demonstrating its ability to avert aversive responses emerging during acute abstinence from opioids. Furthermore, in stress naïve mice, we demonstrated that (*2R,6R*)-HNK reverses maladaptive behavioral responses induced by protracted opioid withdrawal, while in stress-exposed mice, it reversed stress susceptibility induced by prior opioid exposure. Importantly, it also showed robust prophylactic actions against subsequent stress-induced reinstatement of such maladaptive behaviors.

Additionally, we showed for the first time that (*2R,6R*)-HNK is able to facilitate extinction from opioid conditioning and to prevent stress-induced reinstatement of opioid-seeking behaviors. Importantly, (*2R,6R*)-HNK reduced voluntary morphine preference and consumption in mice previously exposed to forced (non-contingent) opioids. The actions of this metabolite to restore maladaptive behavioral responses during protracted opioid withdrawal could stem from its actions to restore synaptic plasticity markers, including hippocampal BDNF, which we found to be reduced during protracted morphine withdrawal. These collective findings highlight the multifaceted potential of (*2R,6R*)-HNK as a therapeutic intervention in opioid addiction, impacting conditioning, acute and protracted withdrawal-related aversive behaviors, stress susceptibility, and synaptic adaptations, thus emphasizing its promise as a comprehensive approach for managing opioid addiction-related behaviors.

The escalating misuse of opioid-based prescription treatments has led to widespread addiction over the last decades, with chronic opioid abuse resulting in persistent negative affective symptoms such as lowered mood, anhedonia, and social avoidance, acting as triggers for relapse during protracted abstinence ^7,9^. Currently available pharmacotherapies have limited efficacy or require life-long administration to prevent relapse and some treatments are opioid-based substitutes, possessing high risk to also be abused ^39^. To effectively treat opioid addiction, there is an urgent need to identify drugs that will rapidly alleviate negative affective symptoms and prevent relapse during protracted opioid abstinence. Ketamine, an NMDAR antagonist recognized for its rapid antidepressant effects, has shown promise in enhancing abstinence rates and decreasing craving and somatic responses to precipitated opioid withdrawal in opioid-dependent individuals ^15–17^. Similar to the clinical findings, racemic ketamine ^18^ and (*R*)-ketamine ^19^ reduced somatic symptoms of precipitated withdrawal from opioids in rodents. Overall, sub-anesthetic doses of ketamine were shown to reduce opioid-seeking behaviors, as well as acute somatic withdrawal symptoms in both clinical and preclinical studies ^17,40–43^. Despite its potential in addressing certain facets of opioid addiction, the practical application of ketamine for treating opioid addiction is restricted due to its dissociative effects and the risk of abuse, even at low doses ^20^.

Ketamine is rapidly and stereoselectively metabolized to several metabolites, including *N*-demethylated norketamines and dehydronorketamines, as well as hydroxynorketamines (HNKs) ^21^. We have previously provided evidence that (*2R,6R*)-HNK, similar to its parent drug ketamine, rapidly reverses negative affective behaviors via a presynaptic mechanism converging with the mGlu_2_ receptor signaling to enhance excitatory neurotransmission, which subsequently activates downstream BDNF-related pathways ^22–27^. There is only one recent preclinical study in mice showing that the (*2R,6R*)-HNK metabolite is able to reduce acute withdrawal precipitated somatic symptoms, and to block priming-induced reinstatement of opioid-seeking behaviors following extinction ^29^. Although these prior findings are promising, they do not provide evidence for the efficacy of (*2R,6R*)-HNK in reversing the negative affect, or stress-induced reinstatement during protracted opioid withdrawal. Here, we demonstrate a significant effectiveness of this metabolite of ketamine to reverse affective symptoms associated with complex interactions between protracted opioid abstinence and stress susceptibility. Therefore, our findings hint possible therapeutic potential of (*2R,6R*)-HNK to treat comorbid opioid use and mood disorders. It is critical to tackle the emotional challenges linked to opioid abstinence due to the substantial overlap with depression, especially considering the uncertainty surrounding the effectiveness of traditional antidepressants in treating this group of patients ^44^.

While often overlooked, extended use of addictive substances frequently leads to social deterioration characterized by isolation and impaired decision-making, prioritizing an obsessive focus on the drug and its associated usually environmental triggers ^45,46^. The demonstrated enhancement of sociability in morphine-withdrawn mice by (*2R,6R*)-HNK holds significant importance, addressing a pivotal emotional disruption that renders patients susceptible to relapse even after prolonged abstinence. This is particularly noteworthy as our study represents one of the initial efforts to rescue sociability deficits linked with opioid addiction. Considering the efficacy of social support programs and the advantages of social reintegration in maintaining abstinence among former addicts ^47^, the utilization of compounds that enhance sociability in individuals abstaining from opioids could potentially be highly effective in averting relapse.

While the mechanisms underlying the efficacy of (*2R,6R*)-HNK in preventing mood disturbances during protracted abstinence from opioids is not clear, it is possible that this compound reverses overall synaptic plasticity impairment in mood-regulating 2GluN2A-NMDARs induced by prolonged opioid abstinence in ventral hippocampal synapses. We, and others, previously showed that BDNF-TrkB signaling is among the most possible drug targets underlying HNK’s pharmacological actions ^22,27,48–50^. Notably, we recently provided evidence suggesting that both ketamine and its biologically active metabolite, (*2R,6R*)-HNK, are likely to exert their unique rapid reduction of negative affective behaviors by activating the GluN2A-NMDARs ^32^. Our findings in the present study, demonstrating for the first time that (*2R,6R*)-HNK can reverse decreased levels of synaptic GluN2A-NMDARs during protracted opioid abstinence, align with a mechanism whereby the actions of (*2R,6R*)-HNK converge on this receptor system. Continuing efforts aimed at unraveling the precise molecular and cellular mechanisms underlying the observed therapeutic efficacy of (*2R,6R*)-HNK in opioid addiction, coupled with exploring its effectiveness in diverse opioid use models, are crucial steps toward understanding the broader potential of this ketamine metabolite as an innovative, next-generation effective pharmacotherapy of opioid addiction.

## Acknowledgements

The authors would like to thank Dr. Christiana Neophytou and Prof. Panagiotis Papageorgis from the European University for their assistance with the Western Blot studies. We would also like to thank Mr. Chris Powels for his contribution in performing the initial scoring of the precipitated withdrawal-related somatic symptoms (Experiment 4). Research was supported by a Brain and Research Foundation (NARSAD; #26826) Young Investigator Grant to P.Z. and Research & Innovation Foundation of Cyprus – Excellence Hubs 2021 (EXCELLENCE/0421/0543) to PZ, with A.O. being listed as a co-I, and a European Commission Marie Skłodowska-Curie fellowship #101031962 to P.Z. and #01031317 to P.G. This publication was made possible by support from the IDSA Foundation. Its contents are solely the responsibility of the authors and do not necessarily represent the official views of the IDSA Foundation.

