## Supplemental Figure 1 for "(*2R,6R*)-hydroxynorketamine facilitates extinction and prevents emotional impairment and stress-induced reinstatement in morphine abstinent mice"

**Affiliations:**
[1] Department of Psychology, University of Cyprus, Nicosia, 2109, Cyprus.
[2] Department of Biology, University of Cyprus, Nicosia, 2109, Cyprus.
[3] Department of Psychology, University of Wisconsin-Milwaukee, USA.
[4] Department of Psychiatry, School of Medicine, University of Maryland, Baltimore, MD, USA.

[*] Equal contribution

✉ **Correspondence to**:

**Dr. Panos Zanos**

Assistant Professor,
Department of Psychology
University of Cyprus
Director, Translational Neuropharmacology Lab
Aglantzia, Nicosia, Cyprus, 2109


### SUPPLEMENTARY FIGURE AND FIGURE LEGEND

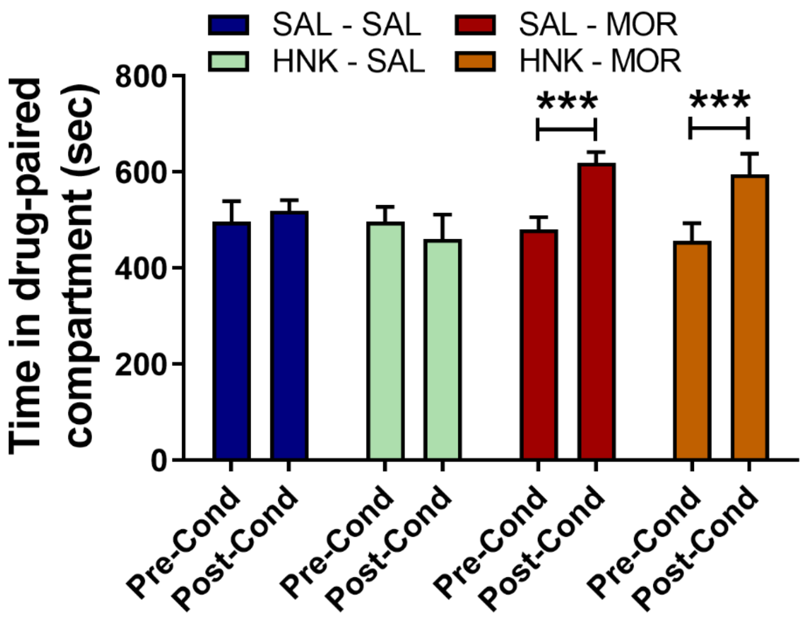


**Supplementary Figure 1. Effect of (*2R,6R*)-HNK on Morphine Conditioning.** We examined the impact of (*2R*,*6R*)-HNK (10 mg/kg) on conditioning to high doses of morphine (5 mg/kg). Administration of (*2R*,*6R*)-HNK did not inhibit the development of high-dose morphine CPP in mice. Data are the mean ± S.E.M.; n= 10-11 mice/group; * $p<0.05$, ** $p<0.01$. *Abbreviations*: Cond, conditioning; HNK, hydroxynorketamine; MOR, morphine; SAL, saline
